# High-dimensional characterization of IL-10 production and IL-10 dependent regulation during primary gammaherpesvirus infection

**DOI:** 10.1101/490789

**Authors:** Abigail K. Kimball, Lauren M. Oko, Rachael E. Kaspar, Linda F. van Dyk, Eric T. Clambey

## Abstract

Interleukin (IL)-10 is a potent immunomodulatory cytokine produced by multiple cell types to restrain immune activation. Many herpesviruses use the IL-10 pathway to facilitate infection, but how endogenous IL-10 is regulated during primary infection in vivo remains poorly characterized. Here, we infected mice with murine gammaherpesvirus 68 (*γ*HV68) and analyzed the production, and genetic contribution, of IL-10 by mass cytometry (cytometry by time-of-flight, CyTOF) analysis. *γ*HV68 infection elicited a breadth of effector CD4 T cells in the lungs of acutely infected mice, including a highly activated effector subset that co-expressed IFN*γ*, TNF*α*, and IL-10. By using IL-10 green fluorescent protein (gfp) transcriptional reporter mice, we identified that IL-10 was primarily expressed within CD4 T cells during acute infection in the lungs. IL10gfp expressing CD4 T cells were highly proliferative and characterized by the expression of multiple co-inhibitory receptors including PD-1 and LAG-3. When we analyzed acute *γ*HV68 infection of IL-10 deficient mice, we found that IL-10 limits the frequency of both myeloid and effector CD4 T cell subsets in the infected lung, with minimal changes at a distant mucosal site. These data emphasize the unique insights that high-dimensional analysis can afford in investigating antiviral immunity, and provide new insights into the breadth, phenotype and function of IL-10 expressing effector CD4 T cells during acute virus infection.

This work was funded by National Institutes of Health Grants R01 AI121300 and R01 CA168558 (to L.F.v.D.), an American Heart Association National Scientist Development grant (#13SDG14510023), the Crohn’s and Colitis Foundation of America (#311295), a pilot grant from the Lung, Head and Neck Cancer program within the University of Colorado Cancer Center, and a Career Enhancement Award from the University of Colorado Lung Cancer Specialized Program of Research Excellence (P50CA58187) (all to E.T.C.).

The Lung, Head and Neck program within the University of Colorado Cancer Center, and the Flow Cytometry Shared Resource, are directly funded through support from the National Cancer Institute Cancer Center Support Grant P30CA046934.

**Abbreviations used in this article:**

B6: C57BL/6
CyTOF: cytometry by time-of-flight
*γ*HV68: murine gammaherpesvirus 68
IFN*γ*: Interferon-gamma
IL-10: Interleukin-10
ICCS: Intracellular cytokine stain
KO: knockout
TNF*α*: tumor necrosis factor-alpha
tSNE: t-distributed stochastic neighbor embedding.

## INTRODUCTION

The gammaherpesviruses (*γ*HVs) are a group of large dsDNA lymphotropic viruses which include the human pathogens Epstein-Barr virus (EBV) and Kaposi’s sarcoma associated herpesvirus (KSHV), and the small animal model murine gammaherpesvirus 68 (*γ*HV68) (1-3). The *γ*HVs establish a lifelong infection in their host, with most infections in immunocompetent hosts asymptomatic. In contrast, immunosuppressed individuals are at a significantly increased risk for the development of a variety of chronic pathologies, including *γ*HV-associated malignancies (4).

*γ*HV infection is critically regulated by the immune system, and *γ*HVs have evolved numerous strategies to either subvert or avoid immune destruction, thereby facilitating lifelong infection (5-7). Among these, IL-10 is a multifunctional cytokine that downregulates the expression of multiple pro-inflammatory cytokines and cell surface molecules, and can influence a wide array of immune cell types (8-10). IL-10 is a frequent target of manipulation by the herpesviruses. Some *γ*HVs, including Epstein-Barr virus (EBV), encode their own IL-10 homolog (11, 12). In contrast, other viruses, including murine gammaherpesvirus 68 (*γ*HV68), inducing cellular IL-10 (13). IL-10 has been reported to regulate multiple aspects of *γ*HV68 infection. For instance, *γ*HV68-infected IL-10 deficient mice fail to control leukocytosis, have greater splenomegaly and a reduced latent load (14), with these mice susceptible to an exacerbated inflammatory bowel disease (15). IL-10 has been reported to be induced in B cells by the *γ*HV68 M2 gene product (13). IL-10 expressing CD8 T cells have also been observed during chronic *γ*HV68 infection in mice depleted for CD4 T cells, an immunosuppressive phenotype associated with chronic viral pathogenesis (16).

Based on the aforementioned studies, IL-10 regulates *γ*HV68 infection. Here we applied high-dimensional, single-cell analysis using mass cytometry (cytometry by time-of-flight, CyTOF) (17) to define the breadth and cellular phenotype of IL-10 expressing cells elicited during acute *γ*HV68 infection. We further analyzed the genetic impact of IL-10 in limiting *γ*HV68 induced inflammation. These studies demonstrate the acute impact of IL-10 on primary *γ*HV infection in vivo and define effector CD4 T cells as a major cell source of IL-10 during acute pulmonary infection.

## MATERIALS AND METHODS

### Experimental samples

Mice were obtained from the Jackson Laboratory and bred in-house at the University of Colorado, including the C57BL/6J (B6, Jax stock #000664), IL10-deficient (IL10KO, B6.129P2-*Il10*^*tm1Cgn*^/J, Jax stock #002251), or IL10gfp (B6.129S6-*Il10*^*tm1Flv*^/J, Jax stock #008379) genotypes. Mice were intranasally infected with 4×10^5^ plaque forming units (PFU) of wild-type (WT) murine gammaherpesvirus 68 (*γ*HV68) in mice subjected to isoflurane induced anesthesia. Mice were used between 8-15 weeks of age, with experimental cohorts age-and sex-matched. Mice were infected with WT *γ*HV68 (strain WUMS, ATCC VR-1465) (18), using either bacterial artificial chromosome (BAC)-derived WT *γ*HV68 (19) or WT *γ*HV68.ORF73*β*la, which encodes a fusion between ORF73 and the beta-lactamase gene (20). *γ*HV68 was grown and titered by plaque assay on NIH 3T12 fibroblasts as previously published (21). Mice subjected to anti-CD3 antibody injection were injected intraperitoneally with 15 μg of anti-CD3*ε* antibody (clone 145-2C11, BioXcell, Cat# BP0001-1) at 0 and 46 hours, with spleens harvested at 50 hours post primary injection (22). Lung data in Figure 4 are from a previously published dataset (23). All procedures were performed under protocols approved by the Institutional Animal Care and Use Committee at the University of Colorado Anschutz Medical Campus.

### Cell processing & antibody staining

In-depth processing and staining protocol can be found in (23). Briefly, lungs were perfused using 10-12 mL of phosphate buffered saline (PBS), harvested, minced and enzymatically digested with collagenase D for 1 h at 37°C. Lungs and spleens were further subjected to mechanical disruption to generate single-cell suspensions that were subjected to red blood cell lysis, and resuspended for staining. Colons were surgically dissected, rinsed then vortexed in PBS to remove fecal material, and incubated with PBS (Gibco) containing 15 mM HEPES (HyClone), 1 mM EDTA (ThermoFisher) at room temperature for 15 minutes while samples were vigorously vortexed, to remove intraepithelial lymphocytes. Colonic tissue was then rinsed over a sieve, washed with ice cold PBS, minced and subjected to enzymatic digestion using Collagenase VIII (100-200 units/mL final concentration, Sigma Aldrich) diluted in RPMI 1640 (Gibco) supplemented with 5% fetal bovine serum, 15 mM HEPES, and 1% penicillin/streptomycin. Enzymatic digestion was done for 20 minutes at 37° C, under continuous vortexing. Enzymatic digestion was quenched using ice cold RPMI 1640 with 5% fetal bovine serum. Colonic samples were washed with a 1:10,000 dilution of Benzonase^®^ Nuclease (≥250 units/μL, Sigma Aldrich) in RPMI 1640 to minimize cell aggregation, prior to staining with cisplatin as a live/dead discriminator. For samples analyzed for intracellular cytokine staining, cells were pharmacologicaly stimulated with PMA and ionomycin (ThermoFisher) in the presence of the Golgi apparatus inhibitors brefeldin A and monensin (Biolegend) for 5 hours. Cells were stained with cisplatin according to manufacturer’s recommendations (Cell-ID^™^ Cisplatin, Fluidigm), incubated with Fc receptor blocking Ab (clone 2.4G2, Tonbo Biosciences) for 10-20 min, stained with primary surface Abs for 30 min at 22°C or 15 min at 37°C and 15 min at 22°C. Secondary surface stains, to detect fluorophore conjugated antibodies, were incubated for 20-30 min, and washed, with intracellular staining done using the FoxP3 Fix/Perm Buffer kit (Thermo Fisher) for 2 hours or overnight at 4°C. Following cell staining, cells were washed and resuspended in Intercalator (Cell-ID^™^ Intercalator-Ir). A subset of experiments (Figures 1, 3, 4B, 4D, 4F, and 5) were subjected to isotopic barcoding using the Fluidigm barcoding kit (Cell-ID^™^ 20-Plex Pd Barcoding Kit, Fluidigm) prior to staining with the primary surface stains. Antibodies used for these studies are listed in Tables 1-4B. All antibodies that were directly conjugated to isotopically purified elements were obtained from Fluidigm. In each of the CyTOF panels, a subset of antibodies were detected using a secondary detection approach, with FITC, PE, APC or Biotin-conjugated antibodies (clone and source identified in Tables 1-4B), detected using metal conjugated secondary antibodies against FITC, PE, APC or Biotin (Fluidigm).

**Table 1.** Antibody conjugates used for the analysis in Figure 1 and 5.

| Tag | Target | Ab Clone | Surface/Intracellular | Clustering* |
| --- | --- | --- | --- | --- |
| <sup>89</sup>Y | CD45 | 30-F11 | Surface | Yes |
| <sup>141</sup>Pr | Gr1 (Ly6C/Ly6G) | RB6-8C5 | Surface | Yes |
| <sup>142</sup>Nd | CD11c | N418 | Surface | Yes |
| <sup>143</sup>Nd | GITR (CD357) | DTA1 | Surface | Yes |
| <sup>144</sup>Nd | IL-2 | JES6-5H4 | Intracellular | Yes |
| <sup>145</sup>Nd | CD69 | H1.2F3 | Surface | Yes |
| <sup>146</sup>Nd | CD8α | 53-6.7 | Surface | Yes |
| <sup>148</sup>Nd | CD11b | M1/70 | Surface | Yes |
| <sup>149</sup>Sm | CD19 | 6D5 | Surface | Yes |
| <sup>150</sup>Nd | CD25 | 3C7 | Surface | Yes |
| <sup>151</sup>Eu | CD64 | X54-5/7.1 | Surface | Yes |
| <sup>152</sup>Sm | CD3ε | 145-2C11 | Surface | No |
| <sup>153</sup>Eu | PD-L1 (CD274) | 10F.9G2 | Surface | Yes |
| <sup>154</sup>Sm | CTLA4 (CD152) | UC10-4B9 | Intracellular | Yes |
| <sup>155</sup>Gd | IRF4 | 3E4 | Intracellular | Yes |
| <sup>156</sup>Gd | FoxP3-PE / anti-PE | FJK-16s (Thermo Fisher) / PE001 (anti-PE) | Intracellular/Secondary | Yes |
| <sup>158</sup>Gd | IL-10 | JES5-16E3 | Intracellular | Yes |
| <sup>159</sup>Tb | PD-1 (CD279) | RMP1-30 | Surface | Yes |
| <sup>160</sup>Gd | GMCSF-FITC / anti-FITC | MP1-22E9 (Pharmingen) / FIT-22 (anti-FITC) | Intracellular/Secondary | Yes |
| <sup>161</sup>Dy | Tbet | 4B10 | Intracellular | Yes |
| <sup>162</sup>Dy | TNFα | MP6-XT22 | Intracellular | No (Fig. 1)<br>Yes (Fig. 5) |
| <sup>163</sup>Dy | Lag3-APC / anti-APC | C9B7W (Biolegend) / APC003 (anti-APC) | Surface/Secondary | Yes |
| <sup>164</sup>Dy | IκBα | L35A5 | Intracellular | Yes |
| <sup>165</sup>Ho | IFNγ | XMG1.2 | Intracellular | No (Fig. 1)<br>Yes (Fig. 5) |
| <sup>166</sup>Er | IL-4 | 11B11 | Intracellular | Yes |
| <sup>167</sup>Er | IL-6 | MP5-20F3 | Intracellular | Yes |
| <sup>168</sup>Er | Ki-67 | B56 | Intracellular | Yes |
| <sup>169</sup>Tm | Ly-6A/E (Sca-1) | D7 | Surface | Yes |
| <sup>170</sup>Er | NK1.1 (CD161) | PK136 | Surface | Yes |
| <sup>171</sup>Yb | CD44 | IM7 | Surface | Yes |
| <sup>172</sup>Yb | CD4 | RM4-5 | Surface | No |
| <sup>173</sup>Yb | CD117 (ckit) | 2B8 | Surface | Yes |
| <sup>174</sup>Yb | MHC class II (IA/IE) | M5/114.15.2 | Surface | No |
| <sup>175</sup>Lu | CD127 | A7R34 | Surface | Yes |
| <sup>176</sup>Yb | ICOS (CD278) | 7E.17G9 | Surface | Yes |
| <sup>195</sup>Pt | Cisplatin | Cell-ID Cisplatin |  | No |
| <sup>191</sup>Ir, <sup>193</sup>Ir | Intercalator | Cell-ID Intercalator-Ir |  | No |
| <sup>140</sup>Ce <sup>151</sup>Eu<br><sup>153</sup>Eu <sup>165</sup>Ho<br><sup>175</sup>Lu | Normalization<br>Beads |  |  | No |

\*Clustering parameters used for Figure 1 and 5 differed as indicated.

**Table 2.** Antibody conjugates used for the analysis in Figure 2.

| Tag | Target | Ab Clone | Surface/Intracellular | Clustering* |
| --- | --- | --- | --- | --- |
| <sup>89</sup>Y | CD45 | 30-F11 | Surface | Yes |
| <sup>141</sup>Pr | Gr1 (Ly6C/Ly6G) | RB6-8C5 | Surface | Yes |
| <sup>142</sup>Nd | CD11c | N418 | Surface | Yes |
| <sup>143</sup>Nd | CD103-Biotin / anti-Biotin | 2E7 (Biolegend) / 1D4C3 (anti-Biotin) | Surface | Yes |
| <sup>144</sup>Nd | MHC class I | 28-14-8 | Surface | Yes |
| <sup>145</sup>Nd | CD69 | H1.2F3 | Surface | Yes |
| <sup>146</sup>Nd | CD8α | 53-6.7 | Surface | Yes |
| <sup>150</sup>Nd | CD27 | LG.3A10 | Surface | Yes |
| <sup>151</sup>Eu | CD25 | 3C7 | Surface | Yes |
| <sup>152</sup>Sm | CD3ε | 145-2C11 | Surface | Yes (A) / No (B-D) |
| <sup>154</sup>Sm | CD11b | M1/70 | Surface | Yes |
| <sup>156</sup>Gd | CD49b-PE / anti-PE | HMa2 (Biolegend) / PE001 (anti-PE) | Surface/Secondary | Yes |
| <sup>159</sup>Tb | PD-1 (CD279) | RMP1-30 | Surface | Yes |
| <sup>160</sup>Gd | KLRG1-FITC / anti-FITC | 2F1 (Thermo Fisher) / FIT-22 (anti-FITC) | Surface/Secondary | Yes |
| <sup>161</sup>Dy | Tbet | 4B10 | Intracellular | Yes |
| <sup>162</sup>Dy | Ly6C | HK1.4 | Surface | Yes |
| <sup>163</sup>Dy | Lag3-APC / anti-APC | C9B7W (Biolegend) / APC003 (anti-APC) | Surface/Secondary | Yes |
| <sup>164</sup>Dy | IkBα | L35A5 | Intracellular | Yes |
| <sup>166</sup>Er | CD19 | 6D5 | Surface | Yes |
| <sup>167</sup>Er | CD150 | TC15-12F12.2 | Surface | Yes |
| <sup>168</sup>Er | Ki-67 | B56 | Intracellular | Yes |
| <sup>169</sup>Tm | IL10gfp | 5F12.4 (anti-GFP) | Intracellular | Yes |
| <sup>170</sup>Er | CD40L | MR1 | Surface | Yes |
| <sup>171</sup>Yb | CD44 | IM7 | Surface | Yes |
| <sup>172</sup>Yb | CD4 | RM4-5 | Surface | Yes (A) / No (B-D) |
| <sup>173</sup>Yb | CD117 (ckit) | 2B8 | Surface | Yes |
| <sup>174</sup>Yb | MHC class II (IA/IE) | M5/114.15.2 | Surface | Yes (A) / No (B-D) |
| <sup>175</sup>Lu | CD127 | A7R34 | Surface | Yes |
| <sup>176</sup>Yb | ICOS (CD278) | 7E.17G9 | Surface | Yes |
| <sup>195</sup>Pt | Cisplatin | Cell-ID Cisplatin |  | No |
| <sup>191</sup>Ir, <sup>193</sup>Ir | Intercalator | Cell-ID Intercalator-Ir |  | No |
| <sup>140</sup>Ce <sup>151</sup>Eu<br><sup>153</sup>Eu <sup>165</sup>Ho<br><sup>175</sup>Lu | Normalization Beads |  |  | No |

\*Clustering parameters used for Figure 2B-D excluded CD3 $\epsilon$ , CD4, and MHC class II.

**Figure 1.**
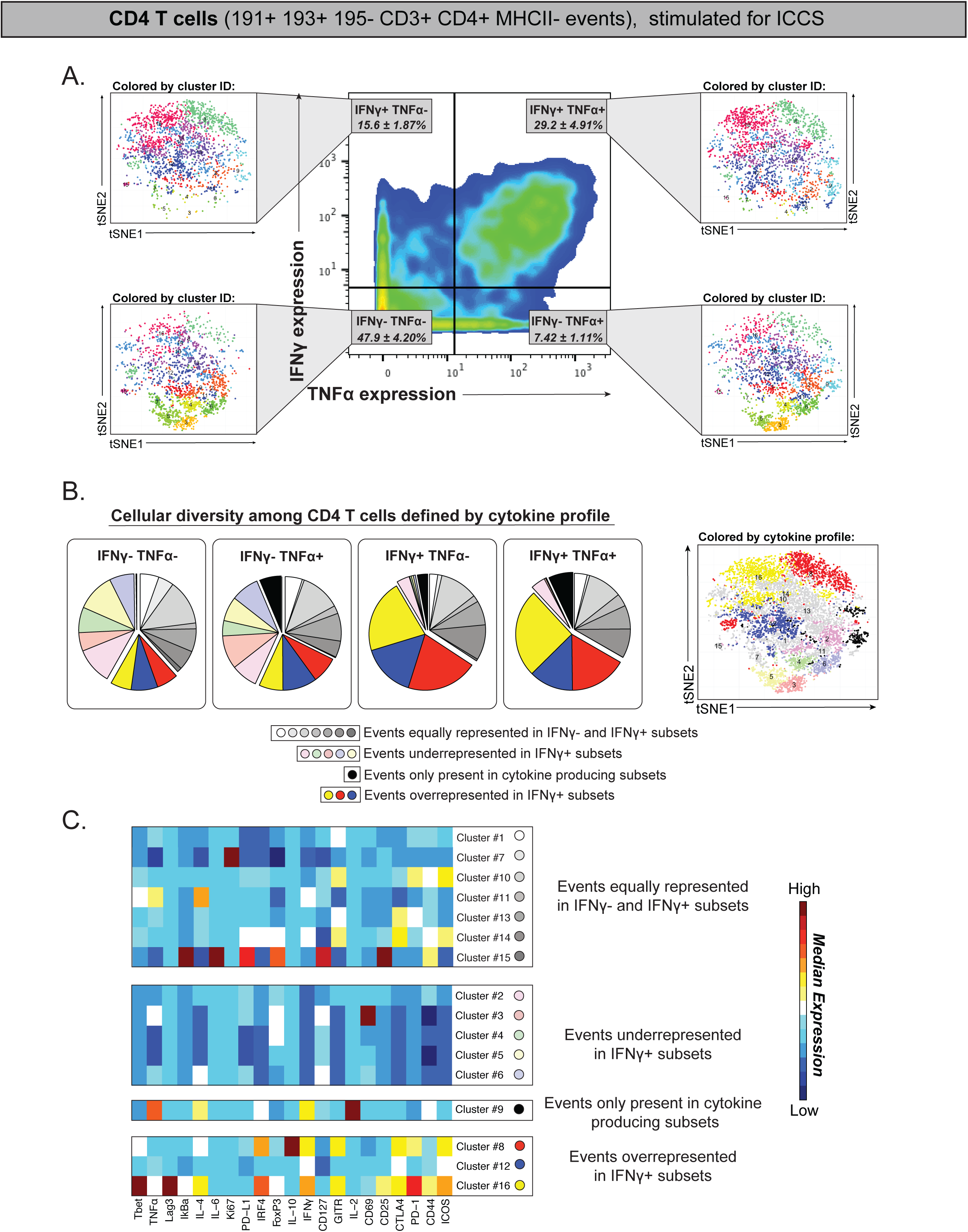
High-dimensional analysis of CD4 T cells elicited during primary γHV68 infection. Mass cytometric analysis of cells recovered from the lungs of *γ*HV68 infected C57BL/6J (B6) mice at 9 dpi, subjected to intracellular cytokine staining (ICCS) analysis, using a 35 antibody panel (Table 1). Data were gated on viable CD4 T cells, defined as ^191^Ir+ ^193^Ir+ ^195^Pt-^152^CD3*ε*+ ^172^CD4+ ^174^MHC II-events, where numbers indicate isotopic mass for each measured parameter. (A) Analysis of IFN*γ* and TNF*α* production from CD4 T cells subjected to pharmacologic stimulation with PMA and ionomycin. The mean ± SEM for each population is identified in each quadrant. Events in each quadrant were further analyzed using the PhenoGraph algorithm and plotted using the tSNE dimensionality reduction algorithm. Events were imported into PhenoGraph and clustered on 8,320 events total and 30 markers (clustering parameters identified in Table 1). In total, 16 clusters were identified, with clusters colored by cluster ID and displayed on tSNE plots. (B) Distribution of PhenoGraph-defined clusters within CD4 T cells, stratified by expression of IFN*γ* and TNF*α*. Each pie chart is subdivided into clusters that are equally represented in IFN*γ*- and IFN*γ*+ CD4 T cells (shades of gray), clusters that are enriched in IFN*γ*-CD4 T cells (pastel colors), clusters that are enriched only in cytokine producing CD4 T cells (black), and clusters that are enriched in IFN*γ*+ CD4 T cells (saturated colors) as defined in the key. Right panel shows events depicted using tSNE, where each cluster is colored according to the cytokine profile subsets (see key). (C) Phenotypic marker expression (in columns) of PhenoGraph-defined clusters (in rows), stratified by their enrichment as a function of cytokine profile. Clusters are stratified as in panel B. Data are from virally infected B6 lungs (n=4 mice) harvested 9 dpi, with cells stimulated with PMA and ionomycin for five hours prior to antibody staining.

**Figure 2.**
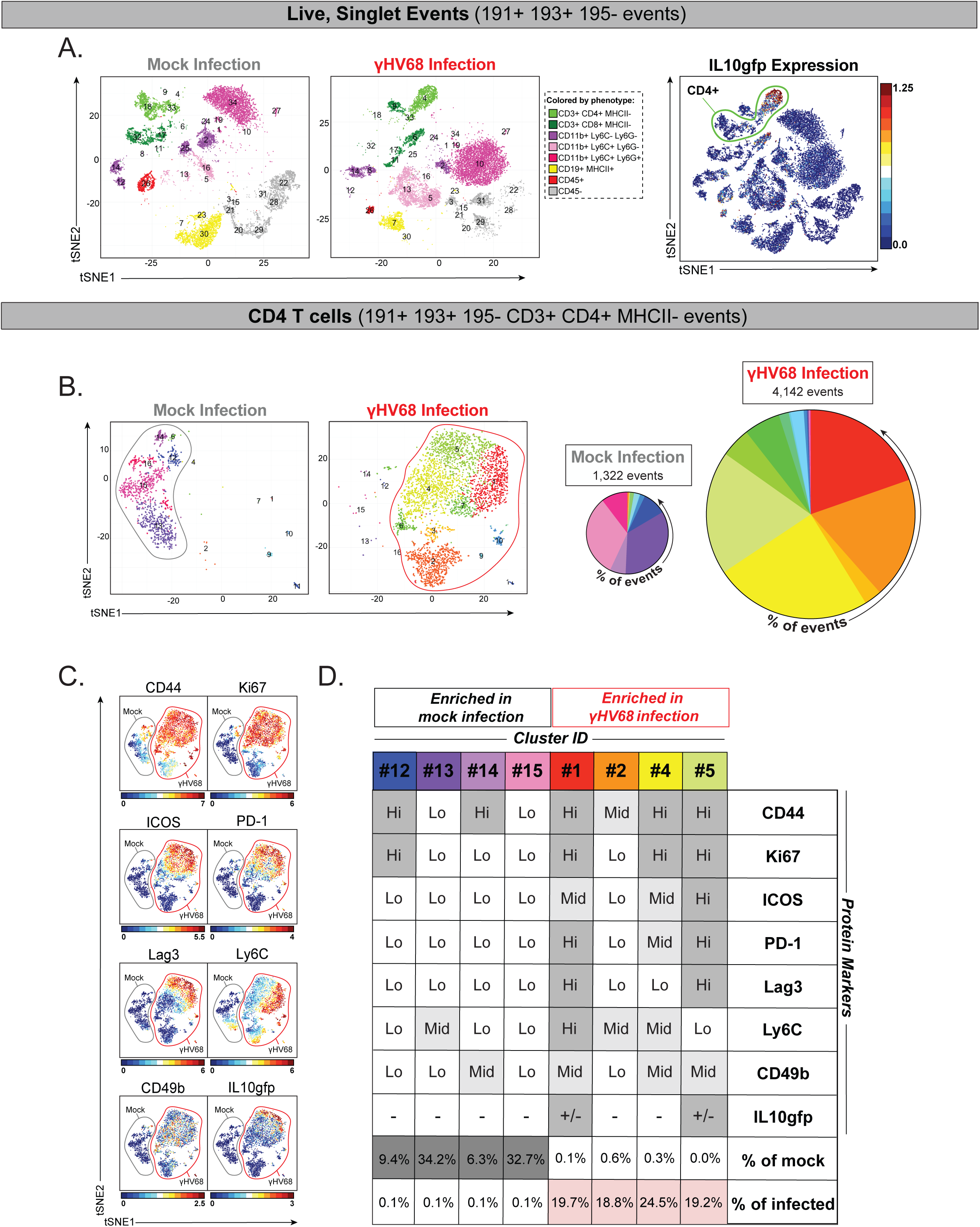
High-dimensional analysis of IL10gfp expression during acute γHV68 infection in the lung. Mass cytometric analysis of cells recovered from the lungs of *γ*HV68 infected IL10gfp mice at 6 dpi. Files were normalized, with events gated on (A) total viable, single cells (defined as ^191^Ir+ ^193^Ir+ ^195^Pt-) or (B-D) viable, CD4+ T cells (defined as ^191^Ir+ ^193^Ir+ ^195^Pt-^152^CD3*ε*+ ^172^CD4+ ^174^MHC II-) prior to analysis, where numbers indicate isotopic mass for each measured parameter. (A) PhenoGraph analysis of cellular phenotypes in mock and virally-infected lung (28,924 events total), clustered based on 29 markers (Table 2), identified 34 unique clusters, with cluster phenotype defined according to the indicated lineage markers. The “CD45+” cluster was defined by its expression of CD45+ and absence of other lineage defining markers. The right panel of A shows all viable, singlet events from mock- and virally-infected lungs, with events colored according to IL10gfp expression. The green boundary line defines CD4+ T cells. (B) Mass cytometric analysis and PhenoGraph-based cell clustering of CD4+ T cells (5,464 events total, clustered based on 26 markers, excluding CD3, CD4, and MHC II, Table 2). In total 16 unique clusters were identified, visualized on a tSNE plot (left). The number and frequency of CD4 T cell clusters is shown in the right panel, with pie charts sized proportionally to the relative cell number of CD4 T cells in mock or virus infected lung. (C) Phenotypic analysis of CD4 T cells, with events predominantly in mock infection denoted with a gray boundary and events predominantly in virus infection denoted with a red boundary. Data depict CD4 T cells plotted according to tSNE1 and tSNE2, as in panel B, with individual plots depicting relative protein expression for the identified marker portrayed by color intensity, with range of expression indicated on the bottom of each panel. Panels are ordered based on the frequency of positive events. (D) Summary of CD4 T cell clusters (in columns) that differ between mock and *γ*HV68 infected lungs, with protein expression denoted in rows (gray shading scaled relative to expression level). Data are from the lungs of a mock- or *γ*HV68-infected mouse harvested at 6 dpi, with panel D quantifying the frequency of events in each cluster as a percentage of CD4+ T cells.

**Figure 3.**
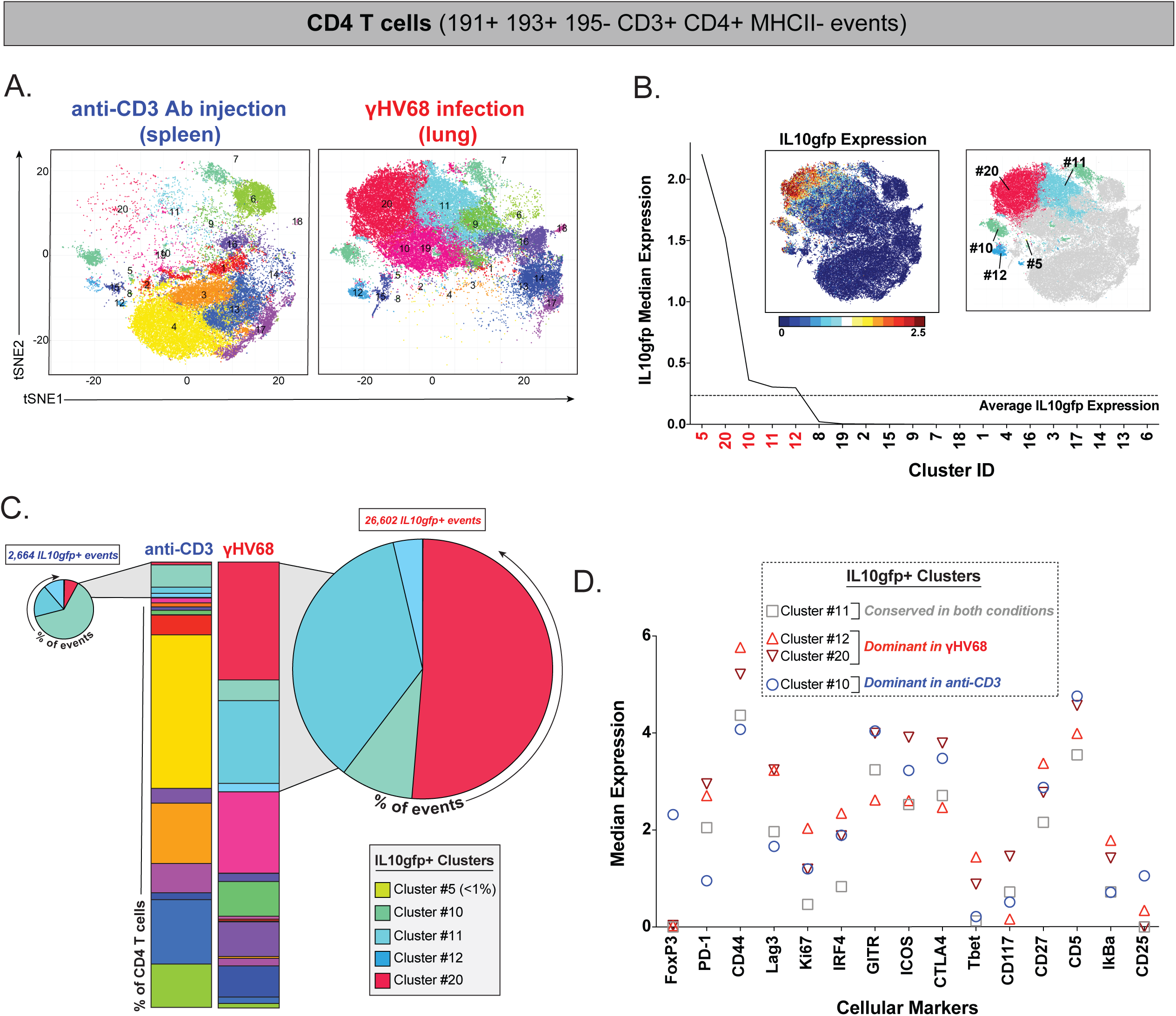
γHV68 infection elicits a dominant IL10gfp expressing CD4 T cell population that differs relative to IL10gfp expressing CD4 T cells in anti-CD3 antibody injected mice. Mass cytometric analysis of cells recovered from IL10gfp mice, comparing CD4 T cell phenotypes between the spleens of anti-CD3 antibody injected mice with the lungs of *γ*HV68 infected mice harvested at 6 dpi. Files were normalized and gated on viable, CD4+ T cells (defined as ^191^Ir+ ^193^Ir+ ^195^Pt-^152^CD3*ε*+ ^172^CD4+ ^209^MHC II-) prior to analysis, where numbers indicate isotopic mass for each measured parameter. (A) PhenoGraph analysis of CD4 T cells, comparing anti-CD3 antibody injected versus *γ*HV68 infected samples (75,483 events total, clustered on 33 markers, excluding CD3, CD4, and MHC II, Table 3), identified 20 unique clusters, each denoted with a distinct color. (B) PhenoGraph-defined CD4 T cell clusters were ranked based on IL10gfp expression, with 5 clusters having higher than average IL10gfp expression identified in red text. Figure insets depict all events from panel A, colored according to (left panel) IL10gfp expression, or (right panel) cluster ID for 5 IL10gfp+ clusters. (C) The frequencies of CD4 T cell clusters are shown for both conditions, with focused analysis on the distribution of IL10gfp+ clusters identified by shaded gray extensions and pie charts on either side. Pie charts denote the frequencies of IL10gfp+ events in each condition, with pie charts sized according to the relative number of IL10gfp+ events present in each condition. (D) Comparison of median protein expression within IL10gfp+ CD4 T cell clusters in either anti-CD3 antibody injected spleens and/or *γ*HV68 infected lungs. Cellular markers are ordered by range in median expression between clusters, from greatest to least in value. Data are from IL10gfp mice, using either spleens from mice that were injected with an anti-CD3 antibody (injected with antibody at 0 and 46 hours, with harvest at 50 hours, n=4 mice) or lungs from *γ*HV68 infected (n=5 mice) harvested at 6 dpi.

**Figure 4.**
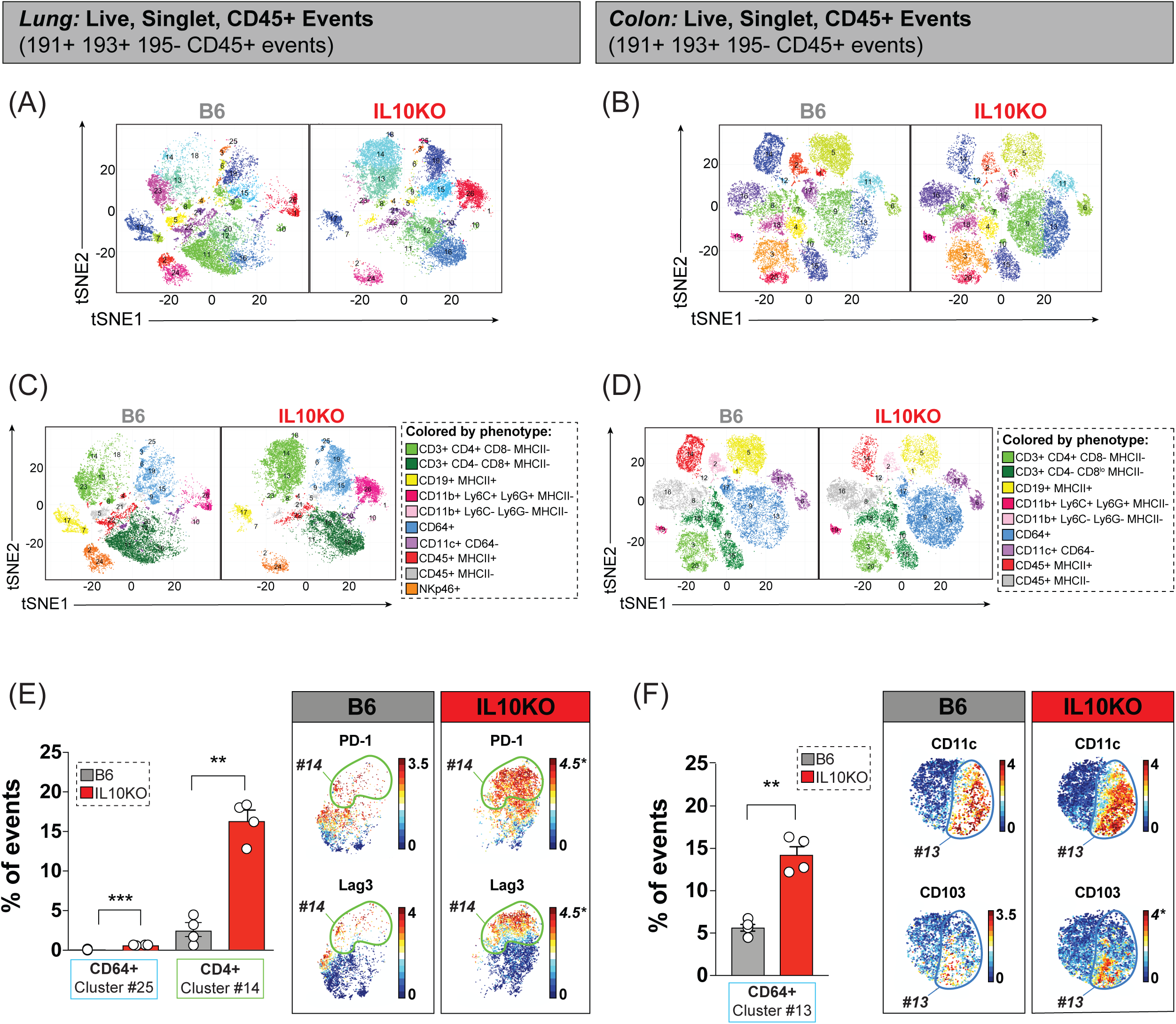
High dimensional analysis of IL-10 dependent regulation of the antiviral response in the lung and the colon. Mass cytometric analysis of cells recovered from lungs (A,C,E) or colons (B,D,F) of *γ*HV68 infected B6 or IL10KO mice harvested at 9 dpi. Files were normalized and gated on viable, single CD45+ cells (defined as ^191^Ir+ ^193^Ir+ ^195^Pt-^89^CD45+) prior to analysis, where numbers indicate isotopic mass for each measured parameter. (A, B) PhenoGraph analysis of CD45+ cells from (A) lungs or (B) colons of *γ*HV68 infected B6 or IL10KO mice. Clustering was done on 4,344 events per file (34,752 total), with clustering in the lung based on 34 markers (29 clusters identified, Table 4A) and clustering in the colon based on 36 markers (20 clusters identified, Table 4B); lungs and colons were harvested from separate cohorts. PhenoGraph-defined clusters are colored according to cluster ID. (C,D) Definition of cellular phenotypes across PhenoGraph-defined clusters, according to the indicated lineage markers, with distinct cell types given unique colors. “CD45+ MHC II+” and “CD45+ MHC II-” clusters were defined by exclusion from other phenotypes. (E,F) Identification of clusters with statistically significant differences in frequency between B6 and IL10KO infected (E) lungs and (F) colons, showing cluster frequencies (left panel) and expression for the identified parameters within the identified cluster (right panel). In the right panel of E, plotted events are CD4+ T cells identified in panel C. In the right panel of F, plotted events are the dominant CD64+ cell clusters identified in panel D. Parameters with a different maximum scale value between B6 and IL10KO are identified by italicized text and an asterisk. Data are from *γ*HV68 infected B6 and IL10KO lungs (n=4 mice per genotype) and colons (n=4 mice per genotype) harvested 9 dpi. Data depict mean ± SEM with individual symbols indicating values from independent samples. All samples were analyzed for statistical significance using unpaired t tests, corrected for multiple comparisons using the Holm-Sidak method, with statistical significance denoted as * p<0.05, ** p<0.01, ***p<0.001.

### CyTOF run and sample normalization/debarcoding

Samples were collected on a Helios mass cytometer (Fluidigm), with samples resuspended with equilibration beads to allow for signal normalization, with the normalization software downloaded from the Nolan laboratory GitHub page (https://github.com/nolanlab) (as in (23)). For experiments where the samples were subjected to isotopic barcoding (Figures 1, 3, 4B, 4D, 4F, and 5) the debarcoding software was used following normalization (https://github.com/nolanlab/single-cell-debarcoder). Normalized, debarcoded data were subjected to traditional Boolean gating in FlowJo, identifying singlets (^191^Iridium (Ir)+ ^193^ Ir+) that were viable (^195^Platinum(Pt)-). These events were then gated and exported for downstream analysis. Additional Boolean gating was later performed for either CD45+ events or CD4+ T cell events, with gating criteria identified within each figure.

**Table 3.** Antibody conjugates used for the analysis in Figure 3.

| Tag | Target | Ab Clone | Surface/Intracellular | Clustering |
| --- | --- | --- | --- | --- |
| <sup>89</sup>Y | CD45 | 30-F11 | Surface | Yes |
| <sup>141</sup>Pr | pSHP2 [Y580] | D66F10 | Intracellular | Yes |
| <sup>142</sup>Nd | CD11c | N418 | Surface | Yes |
| <sup>143</sup>Nd | GITR (CD357) | DTA1 | Surface | Yes |
| <sup>144</sup>Nd | CXCR3-FITC / anti-FITC | CXCR3-173 (Biolegend)/ FIT-22 (anti-FITC) | Surface/Secondary | Yes |
| <sup>145</sup>Nd | CD69 | H1.2F3 | Surface | Yes |
| <sup>146</sup>Nd | CD8α | 53-6.7 | Surface | Yes |
| <sup>147</sup>Sm | pHistone H2A.X (SER139) | JBW301 | Intracellular | Yes |
| <sup>148</sup>Nd | CD11b | M1/70 | Surface | Yes |
| <sup>149</sup>Sm | CD19 | 6D5 | Surface | Yes |
| <sup>150</sup>Nd | CD27 | LG.3A10 | Surface | Yes |
| <sup>151</sup>Eu | CD25 | 3C7 | Surface | Yes |
| <sup>152</sup>Sm | CD3ε | 145-2C11 | Surface | No |
| <sup>153</sup>Eu | PD-L1 (CD274) | 10F.9G2 | Surface | Yes |
| <sup>154</sup>Sm | CTLA4 (CD152) | UC10-4B9 | Intracellular | Yes |
| <sup>155</sup>Gd | IRF4 | 3E4 | Intracellular | Yes |
| <sup>156</sup>Gd | 41BB-PE / anti-PE | 17B5 (Thermo Fisher) / PE001 (anti-PE) | Surface/Secondary | Yes |
| <sup>158</sup>Gd | FoxP3 | FJK-16s | Intracellular | Yes |
| <sup>159</sup>Tb | PD-1 (CD279) | RMP1-30 | Surface | Yes |
| <sup>160</sup>Gd | CD5 | 53-7.3 | Surface | Yes |
| <sup>161</sup>Dy | Tbet | 4B10 | Intracellular | Yes |
| <sup>162</sup>Dy | Tim3 (CD366) | RMT3-23 | Surface | Yes |
| <sup>163</sup>Dy | BCL6 | K112-91 | Intracellular | Yes |
| <sup>164</sup>Dy | IκBα | L35A5 | Intracellular | Yes |
| <sup>165</sup>Ho | Beta-catenin (active) | D13A1 | Intracellular | Yes |
| <sup>166</sup>Er | Arginase-1 | Polyclonal | Intracellular | Yes |
| <sup>167</sup>Er | Gata3 | TWAJ | Intracellular | Yes |
| <sup>168</sup>Er | Ki-67 | B56 | Intracellular | Yes |
| <sup>169</sup>Tm | IL10gfp | 5F12.4 (anti-GFP) | Intracellular | Yes |
| <sup>170</sup>Er | CD49b | HMa2 | Surface | Yes |
| <sup>171</sup>Yb | CD44 | IM7 | Surface | Yes |
| <sup>172</sup>Yb | CD4 | RM4-5 | Surface | No |
| <sup>173</sup>Yb | CD117 (ckit) | 2B8 | Surface | Yes |
| <sup>174</sup>Yb | Lag3 (CD223) | C9B7W | Surface | Yes |
| <sup>175</sup>Lu | CD127 | A7R34 | Surface | Yes |
| <sup>176</sup>Yb | ICOS (CD278) | 7E.17G9 | Surface | Yes |
| <sup>209</sup>Bi | MHC class II (IA/IE) | M5/114.15.2 | Surface | No |
| <sup>195</sup>Pt | Cisplatin | Cell-ID Cisplatin |  | No |
| <sup>191</sup>Ir, <sup>193</sup>Ir | Intercalator | Cell-ID Intercalator-Ir |  | No |
| <sup>140</sup>Ce <sup>151</sup>Eu <sup>153</sup>Eu<br><sup>165</sup>Ho <sup>175</sup>Lu | Normalization Beads |  |  | No |

**Table 4.**
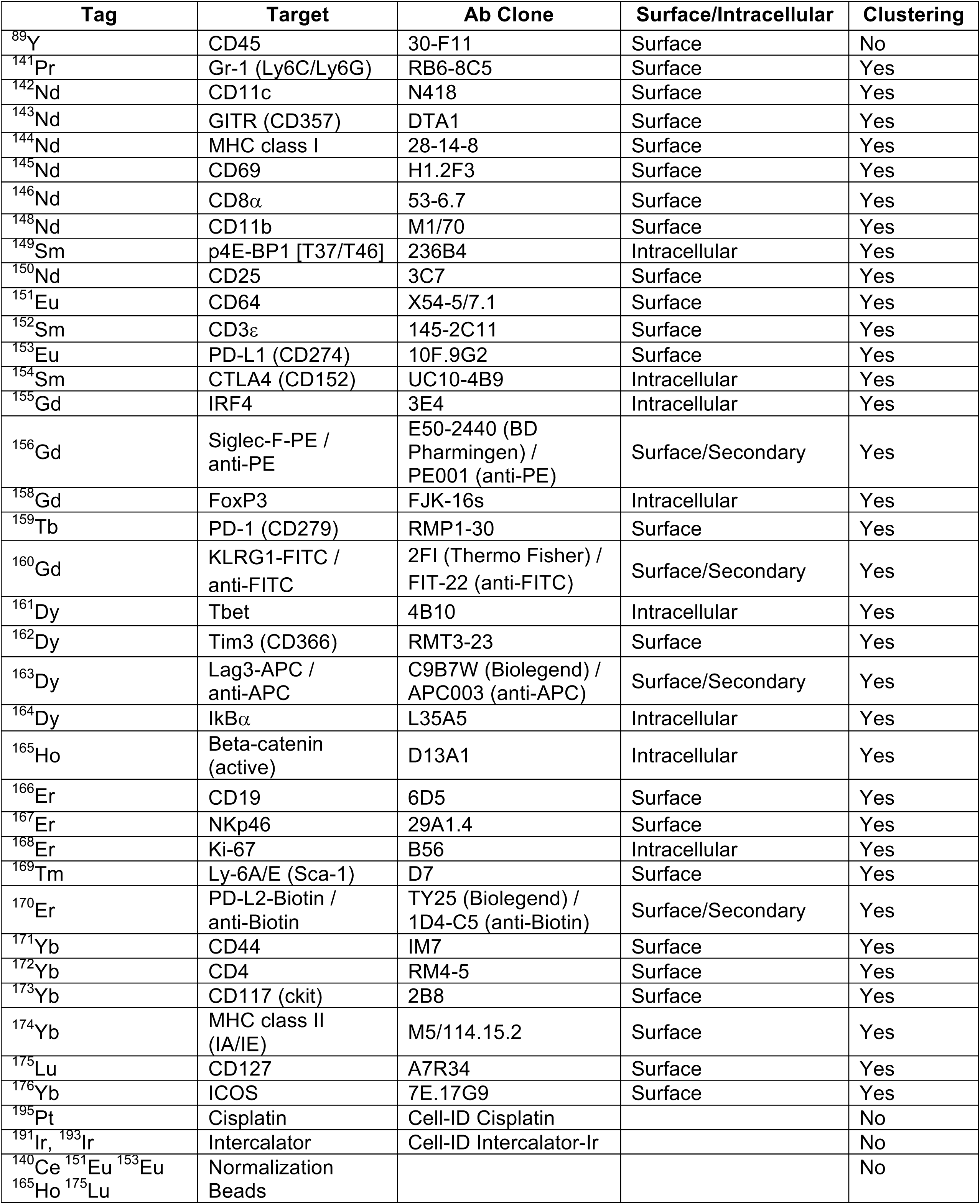
Antibody conjugates used for the analysis in the lung (Figures 4A, 4C, and 4E).

**Table 4B.** Antibody conjugates used for the analysis in the colon (Figures 4B, 4D, and 4F).

| Tag | Target | Ab Clone | Surface/Intracellular | Clustering |
| --- | --- | --- | --- | --- |
| <sup>89</sup>Y | CD45 | 30-F11 | Surface | No |
| <sup>141</sup>Pr | Ly6G | 1A8 | Surface | Yes |
| <sup>142</sup>Nd | CD11c | N418 | Surface | Yes |
| <sup>143</sup>Nd | CD103-Biotin / anti-Biotin | 2E7 (Biolegend) / 1D4C3 (anti-Biotin) | Surface/Secondary | Yes |
| <sup>144</sup>Nd | MHC class I | 28-14-8 | Surface | Yes |
| <sup>145</sup>Nd | SiglecF-PE / anti-PE | E50-2440 (BD Pharmingen) / PE001 (anti-PE) | Surface/Secondary | Yes |
| <sup>146</sup>Nd | CD8α | 53-6.7 | Surface | Yes |
| <sup>148</sup>Nd | CD11b | M1/70 | Surface | Yes |
| <sup>149</sup>Sm | CD19 | 6D5 | Surface | Yes |
| <sup>150</sup>Nd | CD25 | 3C7 | Surface | Yes |
| <sup>151</sup>Eu | CD64 | X54-5/7.1 | Surface | Yes |
| <sup>152</sup>Sm | CD3ε | 145-2C11 | Surface | Yes |
| <sup>153</sup>Eu | PD-L1 (CD274) | 10F.9G2 | Surface | Yes |
| <sup>154</sup>Sm | CTLA4 (CD152) | UC10-4B9 | Intracellular | Yes |
| <sup>155</sup>Gd | IRF4 | 3E4 | Intracellular | Yes |
| <sup>156</sup>Gd | CD90.2 | 30-H12 | Surface | Yes |
| <sup>158</sup>Gd | FoxP3 | FJK-16s | Intracellular | Yes |
| <sup>159</sup>Tb | RORgt | B2D | Intracellular | Yes |
| <sup>160</sup>Gd | CXCR3-FITC / anti-FITC | CXCR3-173 (Biolegend) / FIT-22 (anti-FITC) | Surface/Secondary | Yes |
| <sup>161</sup>Dy | Tbet | 4B10 | Intracellular | Yes |
| <sup>162</sup>Dy | Ly6C | HK1.4 | Surface | Yes |
| <sup>163</sup>Dy | F4/80-APC / anti-APC | MB8 (Thermo Fisher) / APC003 (anti-APC) | Surface/Secondary | Yes |
| <sup>164</sup>Dy | IkBα | L35A5 | Intracellular | Yes |
| <sup>165</sup>Ho | Beta-catenin (active) | D13A1 | Intracellular | Yes |
| <sup>166</sup>Er | Arginase-1 | Polyclonal | Intracellular | Yes |
| <sup>167</sup>Er | NKp46 | 29A1.4 | Surface | Yes |
| <sup>168</sup>Er | Ki-67 | B56 | Intracellular | Yes |
| <sup>169</sup>Tm | Ly-6A/E (Sca-1) | D7 | Surface | Yes |
| <sup>170</sup>Er | CD49b | HMa2 | Surface | Yes |
| <sup>171</sup>Yb | CD44 | IM7 | Surface | Yes |
| <sup>172</sup>Yb | CD4 | RM4-5 | Surface | Yes |
| <sup>173</sup>Yb | CD117 (ckit) | 2B8 | Surface | Yes |
| <sup>174</sup>Yb | Lag3 (CD223) | C9B7W | Surface | Yes |
| <sup>175</sup>Lu | CD127 | A7R34 | Surface | Yes |
| <sup>176</sup>Yb | ICOS (CD278) | 7E.17G9 | Surface | Yes |
| <sup>209</sup>Bi | MHC class II (IA/IE) | M5/114.15.2 | Surface | Yes |
| <sup>195</sup>Pt | Cisplatin | Cell-ID Cisplatin |  | No |
| <sup>191</sup>Ir, <sup>193</sup>Ir | Intercalator | Cell-ID Intercalator-Ir |  | No |
| <sup>140</sup>Ce <sup>151</sup>Eu <sup>153</sup>Eu <sup>165</sup>Ho <sup>175</sup>Lu | Normalization Beads |  |  | No |

### PhenoGraph-based data analysis

Manually gated singlet (^191^Ir+ ^193^Ir+) viable (^195^Pt-) events, or further gated populations, were imported into PhenoGraph, with relevant clustering markers selected (28-35 cellular markers depending on the experiment; note that markers used for gating imported populations were not used for clustering). All parameters used for clustering are indicated in the associated Tables. PhenoGraph was run with the following settings: i) files were merged using either the merge method “min” (Figure 1, Figure 2A, and Figure 3), all (Figure 2B), or “ceiling” (Figure 3-5), ii) files were transformed using the transformation method “cytofAsinh”, iii) the “Rphenograph” clustering method was chosen, coupled with the “tSNE” visualization method. All other settings were automatically chosen, using default PhenoGraph settings.

### PhenoGraph-based visualization

PhenoGraph-defined clusters were displayed on tSNE plots within the R package “Shiny” (23). Within the “Shiny” application, cluster color was altered or colored according to protein expression. Multiple .csv files were produced by PhenoGraph, including “cluster median data” and “cluster cell percentage,” which were used to determine cluster phenotype, distribution between conditions, and statistical significance between groups.

### Software used and statistical analysis

Software for data analysis included: R studio (Version 1.1.453), downloaded from the official R website (http://www.r-project.org/); the Cytofkit package (Version 3.7), downloaded from Bioconductor (https://Bioconductor.org/packages/release/bioc/html/cytofkit.html), Excel 16.15, FlowJo 10.4.2, GraphPad Prism 7.0c, and Adobe Illustrator CC 22.1. Cytofkit was opened using R studio and XQuartz. Statistical significance was tested in GraphPad Prism using an unpaired t test, with statistical significance as identified. For contexts in which we tested statistical significance across all of the identified nodes/clusters (Fig. 4, 5), statistical analysis was subjected to multiple comparison correction in GraphPad Prism.

**Figure 5.**
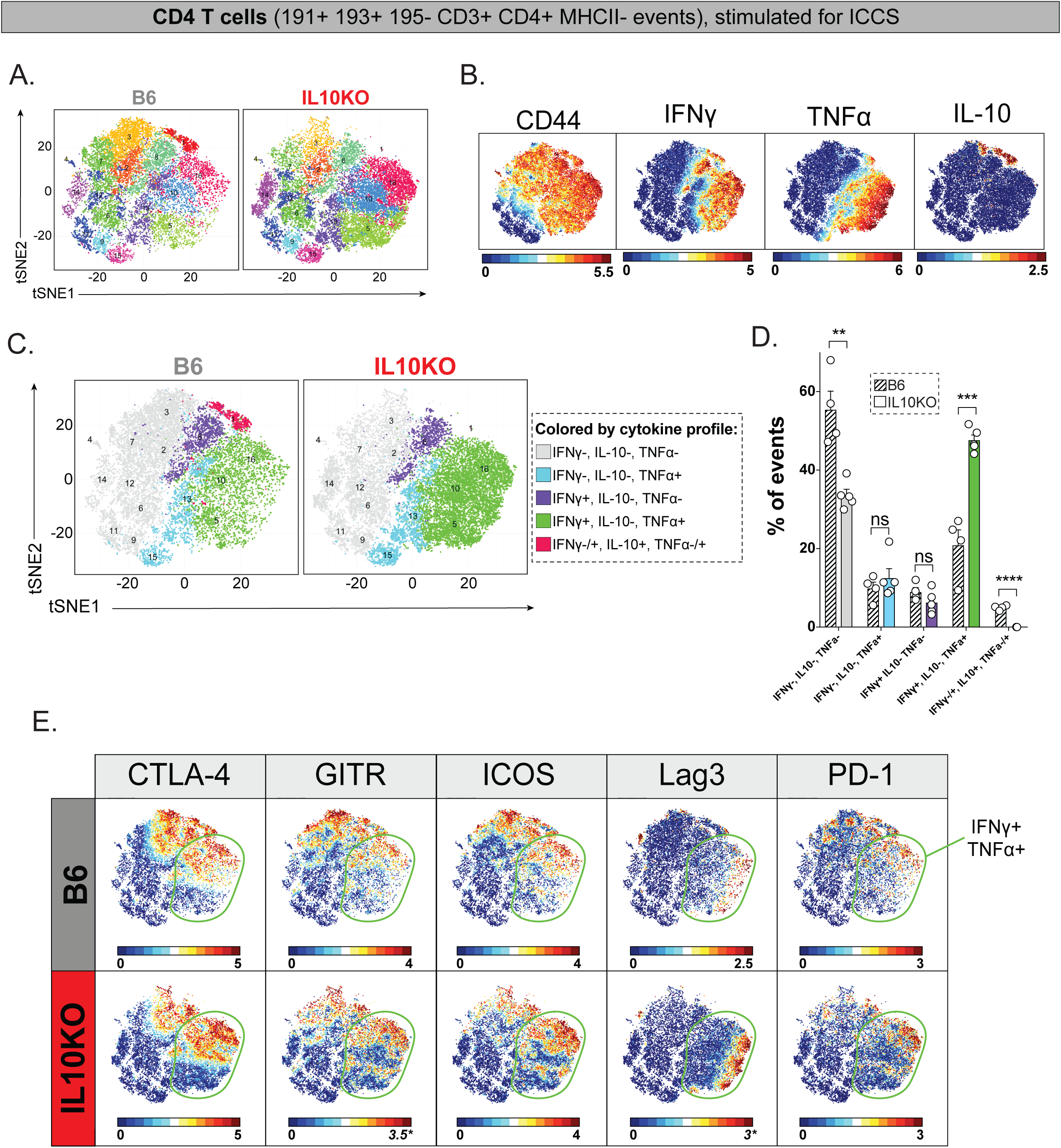
High-dimensional analysis of effector CD4 T cell function following γHV68 infection in B6 and IL10KO mice. Mass cytometric analysis of cells recovered from lungs of *γ*HV68 infected B6 or IL10KO mice harvested at 9 dpi, with cells subjected to pharmacologic stimulation for intracellular cytokine staining analysis (ICCS), using a 35 antibody panel (Table 1). Files were normalized, with data gated on viable CD4 T cells, defined as ^191^Ir+ ^193^Ir+ ^195^Pt-^152^CD3*ε*+ ^172^CD4+ ^174^MHC II-events, where numbers indicate isotopic mass for each measured parameter. (A) CD4 T cells were imported into PhenoGraph and clustered on 33,022 total events (3,669 events from each file) and 32 markers (excluding CD3, CD4, and MHC II, Table 1), identifying 16 unique clusters portrayed on a tSNE plot. (B) All events from panel A are colored by CD44, IFN*γ*, TNF*α*, and IL-10 expression. (C) Events from panel A colored based on their cytokine profile, stratified based on IFN*γ*, TNF*α*, and IL-10 expression. (D) Frequency of CD4 T cells in virally-infected B6 and IL10KO mice. (E) Comparison of CTLA-4, GITR, ICOS, Lag3, and PD-1 expression across CD4 T cells from B6 (top row) or IL10KO (bottom row) mice, where IFN*γ*+ TNF*α*+ CD4 T cells were identified by a green boundary line. Parameters with a different maximum scale value between B6 and IL10KO are identified by italicized text and an asterisk. Data from virally infected lungs of B6 (n=4) and IL10KO (n= 5) mice harvested 9 dpi, with cells stimulated with PMA and ionomycin for five hours prior to ICCS. Data for B6 mice were also included in Figure 1, subjected to different clustering parameters (as outlined in Table 1). Data show mean ± SEM with individual symbols denoting individual mice, with statistical analysis done by unpaired t test, corrected for multiple comparisons using the Holm-Sidak method. Statistical significance denoted by **, p<0.01 and ***, p<0.001. ns, not significant.

## RESULTS

### High-dimensional analysis of effector CD4 T cells during primary γHV68 infection

*γ*HV68 infection induces diverse effector CD4 T cell subsets throughout the course of infection (24). To provide a high-dimensional analysis of effector CD4 T cell function during primary infection, the lungs from *γ*HV68 infected mice were harvested at 9 days post-infection (dpi), subjected to pharmacological stimulation with PMA and ionomycin, and analyzed by mass cytometry using a panel of 35 isotopically purified, metal conjugated antibodies (Table 1). CD4 T cells were initially analyzed for expression of IFN*γ* and TNF*α* expression, two hallmark effector cytokines elicited in antiviral CD4 T cells. Between 40-60% of CD4 T cells harvested from infected lungs expressed either IFN*γ*, TNF*α*, or co-expressed IFN*γ* and TNF*α* (Fig. 1A). IFN*γ*+TNF*α*+ effector CD4 T cells were more frequent than either IFN*γ*+ or TNF*α*+ single positive cells (Fig. 1A). To gain an unbiased perspective on the phenotypic diversity within these cytokine-defined effector CD4 T cells, we next subjected these cell populations to the PhenoGraph algorithm, to define cell clusters present in each cytokine-defined subset (25). By clustering cells based on the expression of 30 proteins (including CD44, Tbet, IRF4 and multiple co-inhibitory receptors, but not on CD3, CD4, MHCII, IFN*γ* or TNF*α*) (Table 1), PhenoGraph defined 16 CD4 T cell clusters present in the virally infected lung (Fig. 1A-B). The relative frequency of these cell clusters was notably affected by whether cells expressed cytokines, and if so, which cytokines. Some clusters, depicted by shades of gray in Fig. 1B, were present in relatively comparable frequencies across all subsets of CD4 T cells, regardless of whether they expressed TNF*α* or IFN*γ* (e.g. cluster #10, a population of CD4 T cells characterized by intermediate expression of GITR, PD-1 and ICOS, Fig. 1C). In contrast, cluster #9 (depicted in black, Fig. 1B-C) contained CD44^high^ IRF4^mid^ IL-2+ cells that were never found among the IFN*γ*-TNF*α*-subset of CD4 T cells. Beyond these distinctions, there were two classes of cell clusters that were inversely related: i) clusters enriched among IFN*γ*-CD4 T cells (depicted in pastel colors), and ii) clusters enriched among IFN*γ*+ CD4 T cells (depicted in saturated colors, Fig. 1B-C). Clusters enriched among IFN*γ*-cells were primarily CD44^low^ with limited expression of notable phenotypic markers (Fig. 1C). Conversely, clusters enriched among IFN*γ*+ subsets (IFN*γ*+ or IFN*γ*+ TNF*α*+; clusters 8, 12 and 16) included: i) cluster #16, a Tbet^high^ Lag3^high^ IRF4^high^ PD-1^high^ GITR+ CD25+ CTLA4+ ICOS+ population and ii) cluster #8, a Tbet^intermediate^ IRF4^high^ IL-10^high^ GITR+ CTLA4+ PD-1+ ICOS+ population (Fig. 1C). These data demonstrate that *γ*HV68 elicits a diverse set of CD4 T cells during primary infection, including the induction of an IFN*γ*+ IL-10+ effector CD4 T cell subset.

### High-dimensional analysis of IL-10 expression as defined by an IL-10 transcriptional reporter

Our initial analysis focused on the phenotypic diversity of CD4 T cells elicited during virus infection, where we identified a prominent IL-10 producing effector CD4 T cell subset (cluster #8, Fig. 1C). To gain a broader perspective on IL-10 expressing cells during primary virus infection, we next infected IL-10 transcriptional reporter mice, in which the enhanced GFP gene is inserted 3’ of the endogenous Il10 gene (i.e. mice expressing the IL-10 *tiger* allele (26)). *γ*HV68 infected lungs were harvested at 6 dpi and subjected to mass cytometric analysis, where samples were stained with a metal-conjugated, anti-GFP antibody to detect the IL-10 reporter (Table 2). As anticipated, *γ*HV68 infection resulted in prominent changes in the frequency and distribution of PhenoGraph-defined clusters relative to mock infected lungs (Fig. 2A). When we analyzed the cellular distribution of the IL10gfp reporter, IL10gfp expression was detected in a fraction of cells in infected lungs, primarily within CD4 T cells (Fig. 2A).

Infection of IL-10 transcriptional reporter mice afforded a major advantage over ICCS, as it measured IL-10 mRNA expression without the need for additional pharmacologic stimuli which can alter cellular phenotype. We therefore focused on CD4 T cell phenotypes in either mock or virally-infected lungs using the PhenoGraph clustering algorithm, which identified 16 CD4 T cell clusters across mock and *γ*HV68 infected lungs. Virally-infected lungs had an increased number of CD4 T cells relative to mock-infection, with a pronounced shift in CD4 T cell clusters (Fig. 2B) towards a prominent CD44^high^ Ki67^high^ ICOS^high^ PD-1^high^ population (Fig. 2C). Within these virally-elicited effector CD4 T cells, Lag3, Ly6C, and CD49b expression were expressed in partially overlapping cell subsets (Fig. 2C). IL10gfp expression was detected in a subset of virally-elicited effector CD4 T cells (Fig. 2C), specifically cluster #1 and #5 (Fig. 2D). IL10gfp+ CD4 T cells between these two clusters shared a conserved CD44^high^ Ki67^high^ ICOS^+^ PD-1^high^ Lag3^high^ CD49b^mid^ phenotype, with Ly6C^high^ and Ly6C^low^ subsets. These studies, using an IL-10 transcriptional reporter, demonstrate that CD4 T cells are a primary source of IL-10 expression during acute *γ*HV68 infection and further identify a core phenotype associated with IL-10 expression within effector CD4 T cells.

### γHV68 infection elicits a dominant IL10gfp expressing CD4 T cell population that differs from anti-CD3 antibody elicited IL10gfp expressing CD4 T cells

Multiple CD4 T cell subsets can produce IL-10, including type 1 regulatory CD4 T cells, a FoxP3-IL-10+ subset of CD4 T cells reported to co-express the cell surface proteins Lag3 and CD49b (22). Based on the expression of Lag3 and CD49b in IL-10 expressing CD4 T cells (Fig. 2C-D), we sought to compare these cells with Tr1 cells generated by an established method. One published method to elicit Tr1 cells is the repeated injection of anti-CD3 antibody into mice, a method associated with both polyclonal T cell activation and the generation of Tr1 cells (22). In this context, Tr1 cells are found especially in the small intestine, with a lower induction of these cells in other tissues (26). To understand how virally induced IL-10+ CD4 T cells compare to IL-10+ CD4 T cells elicited following anti-CD3 antibody injection, we compared CD4 T cells from the lungs of *γ*HV68 infected mice with CD4 T cells from the spleens of mice repeatedly injected with anti-CD3 antibody. Cells were harvested and subjected to mass cytometric analysis using a panel of 34 antibodies (Table 3). When CD4 T cells from these two conditions were subjected to the PhenoGraph algorithm, we identified 20 clusters of CD4 T cells (Fig. 3A). tSNE plots of CD4 T cells for each condition revealed largely non-overlapping cell clusters, suggesting phenotypic divergence (Fig. 3A). Across both conditions, five of twenty CD4 T cell clusters had above average expression of IL10gfp (Fig. 3B). There were large differences in the frequency of IL10gfp+ cells among CD4 T cells between conditions, with ∼10% of CD4 T cells that were IL10gfp+ in the spleens of anti-CD3 antibody injected mice, and ∼50% of CD4 T cells that were IL10gfp+ in *γ*HV68 infected lungs (Fig. 3C). When we analyzed the phenotype of IL10gfp+ clusters between these two conditions, we found a significant phenotypic divergence. Within anti-CD3 antibody injected mice, the highest frequency of IL10gfp+ cells were found within a FoxP3+ regulatory T cell (Treg) cluster (Fig. 3D). In contrast, IL10gfp+ clusters that were dominant in *γ*HV68 infection were CD44^high^ Lag3^high^ PD-1^high^ Tbet+ and FoxP3-, suggesting they may be either Tr1 or Th1 cells (Fig. 3D). These data emphasize the multiple potential cellular sources of IL-10 that can occur with diverse stimuli and across tissues, and reveal that virus infection elicits a distinct effector T cell phenotype when compared to anti-CD3 antibody injection.

### IL-10 dependent regulation of γHV68 infection

To define how IL-10 regulates the distribution and phenotype of immune cell subsets during *γ*HV68 infection we used mass cytometry to compare cellular composition and phenotype between *γ*HV68 infected wild-type (B6) and IL-10 deficient (IL10KO) mice. This analysis focused on cellular diversity among hematopoietic (CD45+) cells within the lungs and colon, with tissues harvested from separate cohorts. Mass cytometry data were collected and subjected to the PhenoGraph algorithm for clustering analysis. This analysis identified 29 cell clusters in lungs harvested from *γ*HV68 infected mice, with 20 cell clusters identified in the colons of *γ*HV68 infected mice (Fig. 4A-B). Cell clusters were further defined based on canonical lineage markers, to quantify the frequencies of distinct leukocyte populations (Fig. 4C-D). When we compared cluster distribution between B6 and IL10KO mice, we found pronounced shifts in cellular distribution between B6 and IL10KO infected lungs, particularly among CD4 T cells (Fig. 4C). In contrast, colons from virally-infected mice had a very limited number of changes in cell clusters at this time post-infection (Fig. 4D). Among the cell clusters present in infected lungs, IL10KO mice had a significantly increased frequency in two cell clusters: i) PD-1+ Lag3+ CD4 T cells (cluster #14), and ii) a small, but significant, increase in CD64+ cells (cluster #25) (Fig. 4E). Colons from infected IL10KO mice had a selective increase in the frequency of CD64+ cells characterized by a CD11b+ CD11c+ phenotype with variable CD103 expression (cluster #13) (Fig. 4F). Cluster #13 was unique among CD64+ clusters in CD11c and CD103 expression relative to other CD64+ clusters in the colon (Fig. 4F). These findings suggest that during acute *γ*HV68 infection, IL-10 constrains the expansion and/or survival of effector CD4 T cells in the lung, and further constrains the frequency of CD64+ mononuclear phagocytic cells in both the lung and the colon.

### IL-10 dependent regulation of effector CD4 T cell function during acute γHV68 infection

Next, we sought to define how IL-10 regulates CD4 T cell effector function during acute *γ*HV68 infection, as revealed by intracellular cytokine staining analysis. We compared CD4 T cell function between *γ*HV68 infected B6 and IL10KO mice by ICCS using mass cytometry. In contrast to the clustering analysis done in Fig. 1, CD4 T cells for both genotypes were subjected to PhenoGraph-defined clustering using 32 markers including IFN*γ*, TNF*α* and IL-10 (Table 1), identifying 16 unique clusters (Fig. 5A). CD4 T cells were predominantly CD44^high^, consistent with a large effector CD4 T cell population in virally-infected lungs at this time (Fig. 5B). 7 of the 16 PhenoGraph-defined clusters showed expression of either IFN*γ*, TNF*α* and/or IL-10 (Fig. 5B), with phenotypes ranging from single expression of IFN*γ*+ or TNF*α*+, coexpression of IFN*γ*+ and TNF*α*+, and triple expression of IFN*γ*+ TNF*α*+ and IL-10+ (Fig. 5B-C). When we analyzed the frequency of CD4 T cells stratified by cytokine expression, we found that B6 and IL10KO mice had comparable frequencies of TNF*α*+ single positive and IFN*γ*+ single positive CD4 T cells (Fig. 5C-D). As anticipated, IL10KO mice had no detectable IL-10+ CD4 T cells (Fig. 5C-D). B6 mice had a significantly increased frequency of cytokine negative CD4 T cells (IFN*γ*-TNF*α*- IL-10-) relative to IL10KO mice (Fig. 5C-D). In contrast, IL10KO mice had a significantly increased frequency of IFN*γ*+ TNF*α*+ effector CD4 T cells (Fig. 5C-D). While IL10KO mice had an increased frequency of IFN*γ*+ TNF*α*+ CD4 T cells relative to B6 mice, both genotypes had phenotypic diversity within this cytokine producing subset, including cell subsets with partially overlapping expression of CTLA-4, GITR, ICOS, Lag3 and PD-1 (Fig. 5E). These data demonstrate that IL-10 constrains the magnitude of highly activated IFN*γ*+ TNF*α*+ effector CD4 T cells during acute *γ*HV68 infection, a population characterized by heterogeneous expression of CTLA-4, GITR, ICOS, Lag3 and PD-1.

## DISCUSSION

IL-10 is a multifunctional cytokine that critically shapes the magnitude and activation status of the immune system in response to infection. In addition to its host immunomodulatory functions, IL-10 is a frequent target of viral manipulation by the herpesviruses (12, 13). Here we sought to investigate how *γ*HV68, a small animal model of gammaherpesvirus infection, intersects with IL-10, both in terms of what cells produce IL-10 and how the overall immune response is influenced by IL-10 during acute infection. For these studies, we have focused on acute, primary infection with *γ*HV68, seeking new insights through the use of high-dimensional mass cytometry (CyTOF) analyses.

IL-10 is known to be produced by a large number of cell types, including CD4 and CD8 T cell subsets and B cells (8). Here we make use of IL-10eGFP transcriptional reporter mice, and direct intracellular staining for IL-10 protein, to identify CD4 T cells as a primary source of IL-10 production during acute *γ*HV68 infection in the lung. IL-10+ CD4 T cells elicited during viral infection were associated with a highly activated effector phenotype, characterized by high expression of CD44 with co-expression of the cytokines IFN*γ* and TNF*α*. IL10gfp+ CD4 T cells were proliferating and characterized by expression of PD-1, Lag3, ICOS, CD49b with variable expression of Ly6C. By querying these cellular phenotypes using mass cytometry, we have further analyzed IL10gfp expression across a wide range of leukocyte subsets. These studies demonstrated focused expression of IL-10 within CD4 T cells in the infected lung with minimal IL-10 expression in other cell subsets at this time.

The expression of IL-10 within this highly activated effector CD4 T cells raises the question of what effector subset(s) express IL-10 in this context. Our data demonstrate that FoxP3+ Tregs are not a prominent source of IL-10 during acute *γ*HV68 infection in the lung. Instead, IL-10+ CD4 T cells appear to be either: i) type 1 regulatory T cells, an IL-10 expressing, FoxP3 negative subset of CD4 T cells frequently characterized by co-expression of CD49b and Lag3 (22), or ii) an IL-10 expressing Th1 subset (27-29). While the IL-10 expressing effector CD4 T cells that we identified here co-express CD49b and Lag3, a proposed marker of Tr1 cells (22), recent studies have emphasized that co-expression of CD49b and Lag3 is not a definitive marker of Tr1 cells (30, 31). Conversely, IFN*γ*+ Th1 cells have been reported to express IL-10 in a variety of settings (32), and *γ*HV68 induced IL-10+ effectors express Tbet, the canonical Th1 transcription factor. Despite these observations, at this time there remains no definitive marker that discriminates between Tr1 and Th1 cells. Alternatively, these IFN*γ*+ IL-10+ expressing cells may represent a distinct effector CD4 T cell subset (33). Recently, Eomesodermin was identified as a transcriptional regulator for IL-10+ effector CD4 T cells, in both Tr1 cells (30) and a potentially distinct IFN*γ*+ IL-10+ effector CD4 T cell (34). Whether IL-10+ CD4 T cells elicited during *γ*HV68 infection express, and require, Eomesodermin remains to be tested.

IL-10 regulates its effects through signals transduced by the heterodimeric IL-10 receptor, targeting multiple cell types (9). High-dimensional analysis of the immune response after *γ*HV68 infection identified a wide spectrum of cells in infected lung and colon, a phenotypic diversity readily characterized through use of the PhenoGraph clustering algorithm and visualized by the tSNE data dimensionality method. When we compared cellular and phenotypic diversity between B6 and IL10KO mice, we found a pronounced increase in the frequency of a PD-1+ Lag3+ CD4 T cell subset and an exaggerated induction of IFN*γ*+ TNF*α*+ effector CD4 T cells. We further found evidence for changes in CD64+ populations in both the infected lung and colon. While IL-10 may directly regulate effector CD4 T cell function (35, 36), CD4 T cell differentiation and effector function may also occur due to altered myeloid function(s) (8, 10, 37, 38). Future studies using targeted disruption of IL-10 signaling in myeloid cells (e.g. macrophages, dendritic cells) and T cells will be required to determine whether IL-10 directly or indirectly modulates effector CD4 T cell differentiation during *γ*HV68 infection. Regardless of the molecular basis of this phenoype, the increased frequency of PD-1+ Lag3+ effector CD4 T cells in IL-10 deficient mice suggests that co-inhibitory receptor expression may function as a compensatory mechanism to constrain pathogenic CD4 T cell function in the absence of IL-10.

Beyond insights on IL-10 dependent regulation during *γ*HV68 infection, these studies demonstrate the power of high-dimensional approaches, such as mass cytometry, to investigate the regulation of the immune response at a global level. By applying mass cytometry to CD4 T cells, our studies provide direct evidence for extensive phenotypic diversity of antiviral effector CD4 T cells elicited during primary viral infection. We anticipate that combining mass cytometry-based studies with host and viral genetics will afford new insights into the underpinnings of antiviral CD4 T cell function.

## ACKNOWLEDGEMENTS

The authors acknowledge Melissa Ledezma for technical assistance, Kristina Terrell, Christine Childs, and Karen Helm for technical support for CyTOF studies including machine operation and management of the CU CyTOF antibody bank, and the CyTOF User Group at the University of Colorado Anschutz Medical Campus for their ongoing collaboration and insights.

## AUTHOR CONTRIBUTIONS

E.T.C. and L.F.v.D. conceived and obtained funding for the studies. L.M.O. and R.E.K. performed all experiments, with input on CyTOF experimental design from A.K.K. and E.T.C. A.K.K. performed all data analysis, data visualization, and wrote the initial manuscript draft with input from E.T.C. A.K.K., L.F.v.D. and E.T.C. wrote the manuscript with input from all authors.

